# FUS/TLS undergoes calcium-mediated nuclear egress during excitotoxic stress and is required for Gria2 mRNA processing

**DOI:** 10.1101/386870

**Authors:** Maeve Tischbein, Desiree M. Baron, Yen-Chen Lin, Katherine V. Gall, John E. Landers, Claudia Fallini, Daryl A. Bosco

## Abstract

Excitotoxic levels of glutamate represent a physiological stress that is strongly linked to amyotrophic lateral sclerosis (ALS) and other neurological disorders. Emerging evidence indicates a role for neurodegenerative disease linked RNA-binding proteins (RBPs) in the cellular stress response. However, the relationships between excitotoxicity, RBP function and pathology have not been explored. Here, we found that excitotoxicity induced the translocation of select ALS-linked RBPs from the nucleus to the cytoplasm within neurons. RBPs affected by excitotoxicity include TAR DNA-binding protein 43 (TDP-43) and, most robustly, fused in sarcoma/translocated in liposarcoma (FUS/TLS). FUS translocation occurs through a calcium-dependent mechanism and coincides with striking alterations in nucleocytoplasmic transport. Further, glutamate-induced upregulation of Gria2 in neurons was dependent on FUS expression, consistent with a functional role for FUS under excitotoxic stress. These findings reveal a link between prominent factors in neurodegenerative disease, namely excitotoxicity, disease-associated RBPs and nucleocytoplasmic transport.

## Introduction

Glutamate is the major excitatory neurotransmitter in the central nervous system. Upon release from pre-synaptic terminals, relatively low levels of glutamate activate metabotropic glutamate receptors as well as the ionotropic receptors: α-amino-3-hydroxy-5-methyl-4-isoxazolepropionic acid (AMPA), N-methyl-D-aspartate and kainite, for normal neurotransmission. However, excessive glutamate exposure overstimulates neurons. This causes a massive influx of calcium, which triggers an excitotoxic cascade involving oxidative damage as well as mitochondrial and ER dysfunction^1^. Excitotoxicity has been implicated in neuronal death and degeneration for various neurological conditions, including the fatal neurodegenerative disease amyotrophic lateral sclerosis (ALS)^1–3^. Pathological evidence for excitotoxity includes elevated levels of glutamate in patient cerebrospinal fluid^4,5^ as well as aberrant processing of the AMPA subunit that controls calcium influx at both the transcript (Gria2) and protein (Glutamate Receptor 2; GluR2) level in patient tissue and disease models^6–8^. Further, ALS-causing mutations are present in D-amino acid oxidase, an enzyme that regulates the degradation of the N-methyl-D-aspartate co-agonist, D-serine^9^. Riluzole, the first FDA approved treatment for ALS, is thought to reduce glutamate signaling through anti-excitotoxic effects^10^. Despite this wealth of knowledge and profound disease relevance, the biological mechanisms underlying the cellular response to excitotoxicity have not been fully elucidated.

RNA-binding proteins (RBPs) have emerged as relevant factors in neurodegenerative disease pathogenesis, particularly in the context of ALS and the related disorder, frontotemporal dementia (FTD)^11^. RBPs belong to a unique class of biomolecules that undergo nucleocytoplasmic shuttling in response to various stimuli, including stress. For instance, the disease-linked RBPs fused in sarcoma/translocated in liposarcoma (FUS/TLS or FUS), TAR DNA-binding protein 43 (TPD-43) and heterogeneous nuclear ribonucleoprotein A1 (hnRNPA1) all exhibit nuclear egress during hyperosmotic stress^12–14^. The purpose of this translocation is unclear, and may represent a functional response to cellular stress^12,15^. In support of this notion, cell viability under hyperosmotic stress is compromised when FUS expression is reduced^12^. However, cell stress also represents a non-genetic factor that likely contributes to neurodegenerative disease pathogenesis^15–17^. Indeed, chronic stress may contribute to the pathological cytoplasmic accumulation of TDP-43 and FUS, prevalent features of ALS and FTD^16–20^. For example, TDP-43 partitions into the insoluble fraction of cultured cells following oxidative stress or heat shock^21,22^ and disease-linked RBPs have been found to aggregate *in vivo* following cerebral ischemia^23^. Intriguingly, the effects of stress on RBP translocation appear selective. While ER stress, oxidative stress and heat shock induce the cytoplasmic accumulation of TDP-43 and other RBPs^24,25^, these stressors fail to elicit a response of FUS^12,26^. Given the physiological relevance of excitotoxicity to neurodegenerative disease, an important but unanswered question is whether excitotoxic stress elicits a functional and/or pathological response from disease-associated RBPs.

Here, we demonstrate that excitotoxic levels of glutamate induce the nuclear egress of several ALS-and FTD-linked RBPs, including FUS, TDP-43 and hnRNPA1 into the cytoplasm of neurons. The nucleocytoplasmic equilibrium of FUS was especially sensitive to excitotoxic stress, as FUS was found to rapidly and robustly accumulate within soma and dendrites of cortical and motor neurons under stress. Further, a glutamate-induced increase in dendritic Gria2 was dependent on FUS, consistent with a role for FUS in glutamatergic signaling during the cellular response to excitotoxic stress. Our results also revealed potentially adverse consequences of excitotoxicity, including the translocation of ALS-linked FUS variants and early signs of nucleocytoplasmic transport dysregulation. This study therefore demonstrates that excitotoxicity can trigger neurodegenerative disease-associated pathologies including cytoplasmic RBP accumulation and nucleocytoplasmic transport decline.

## Methods

### Cell Culture and Stress Application

HEK293-T cells were cultured as described^12^. Dissociated primary cortical neuron cultures were prepared using cortices from C57BL/6 embryonic day 14-15 mice. Embryos were isolated in ice-cold Hanks Buffered Saline Solution (Corning 21-023-CV, Corning, NY, USA) and the meninges removed. Cells were dissociated for 12 minutes in 0.05% Trypsin (Invitrogen 25300-054, Carlsbad, CA, USA) at 37°C, diluted in Dulbecco’s Modified Eagle Medium (Invitrogen 11965118) containing 10% Fetal Bovine Serum (MilliporeSigma F4135, Burlington, MA, USA) and strained with a cell strainer before gently pelleting. Cells were then resuspended in Neurobasal media (Invitrogen 21103049), supplemented with 1% Glutamax (Invitrogen 35050-061), 1% Pen-strep (Invitrogen 15140122) and 2% B-27 (Invitrogen 0080085-SA), and plated at 1.8-2×10^5^ cells/mL on poly-ornithine (final concentration of 1.5 μg/mL; MilliporeSigma P4957) coated plates or coverslips. Neuronal cultures were grown under standard culture conditions (37°C, 5% CO2/95% air) fed every 3-4 days by adding half volumes of supplemented neurobasal media to each well/dish, with additional half changes of media occurring every other feeding. Unless indicated, during the first feeding (DIV 2 or 3) neuron cultures were also treated with a final concentration of 0.5-1µM Cytosine β-D-arabinofuranoside hydrochloride (MilliporeSigma C6645) to inhibit non-neuronal cell growth. Experiments were performed on day *in vitro* (DIV) 14-16.

Primary motor neurons were isolated from embryonic day 12.5 murine spinal cords as described^68^. Briefly, after dissociation in 0.1% trypsin (Worthington LS003707, Columbus, OH, USA) at 37°C for 12 minutes, primary motor neurons were purified using a 6% Optiprep (MilliporeSigma D1556) density gradient and plated on glass coverslips coated with 0.5g/L poly-ornithine and natural mouse laminin (Thermo Fisher 23017015, Waltham, MA, USA). Cells were grown in glia-conditioned Neurobasal medium and supplemented with 2% B27, 2% horse serum (MilliporeSigma H1270), and 10ng/ml BDNF (PeproTech 450-02, Rocky Hill, NJ, USA), GDNF (PeproTech 450-44), and CNTF (PeproTech 450-50). Primary motor neurons were treated on DIV 6-8 with Ionomycin or dimethyl sulfoxide and on DIV 8 with kainic acid. For glutamate experiments, 100mM glutamate (MilliporeSigma G5889) was freshly prepared in neurobasal media and diluted using primary neuron cultured media to achieve 0.1-10µM solutions. To apply stress, neuronal media was replaced with glutamate-containing primary cultured media or primary cultured media alone (glutamate-free control) for 10 minutes. After 10 minutes, treatment media was replaced with primary cultured media for 30 minutes or longer depending on the experiment prior to fixation or lysate collection. Kainic acid (Abcam ab144490) was diluted from 10 mM/ml to 300µM/ml in primary cultured media and added to motor neurons for 10 minutes followed by a replacement with glia-conditioned media for one hour. Stock solutions of 5mM Ionomycin (MilliporeSigma I9657) or 1M sodium arsenite (MilliporeSigma 71287) prepared in prepared in dimethyl sulfoxide (Corning 25-950-CQC) or water were diluted to 10µM or 1 mM in primary cultured media, respectively prior to addition to neurons for one hour. Sorbitol (MilliporeSigma S6021) was directly dissolved in primary cultured media to obtain a final concentration of 0.4M and applied to cells for one hour. For experiments in which ethylene glycol tetraacetic acid (EGTA; MilliporeSigma E3889) was added, a 100mM stock was prepared in water, diluted to 2mM in primary cultured media, and allowed to incubate for 30 minutes prior to use during the experimental time course. Translation was inhibited with 2µM cycloheximide (MilliporeSigma C7698). Neurons were treated with 500nm KPT-330 (Cayman Chemical 18127) dissolved in water on DIV 13 for 48 hours prior to treatment with glutamate as well as during the experimental time course.

### Plasmids and Cloning

Human cDNA for FLAG-HA-tagged wildtype, H517Q, R521G or R495X FUS were cloned into the lentiviral vector, pLenti-CMV-TO-Puro-DEST (Addgene 670-1, Cambridge, MA, USA) using the In-Fusion HD Cloning Plus kit (Clontech 638909, Mountain View, CA). To achieve FUS knockdown, shRNA sequences^48^ were packaged using In-Fusion HD cloning into the lentiviral backbone, CSCGW2 (a generous gift courtesy of Miguel Esteves), which contains a green fluorescent protein (GFP)-reporter expressed under a separate CMV promoter. The shRNA targeting sequences were: 5’-GCAACAAAGCTACGGACAA-3’ (shFUS1) and 5’- GAGTGGAGGTTATGGTCAA-3’ (shFUS2) as well as the scrambled control sequence, 5’- AATTCTCCGAACGTGTCACGT-3’ (shSC). The shuttling reporter, NLS-tdTomato-NES (a generous gift courtesy of Martin Hetzer^33^) was cloned into the pLenti-CMV-TO-Puro-DEST vector backbone (Addgene 670-1) using Gateway BP and LR Clonase reactions (Invitrogen 11789020 and 11791020, respectively). The shuttling reporter contained an NLS sequence (PPKKKRKVQ) and NES sequence (LQLPPLERLTL) attached to tdTomato by a GGGG linker at the N and C termini, respectively.

### Transient Expression of ALS-Mutant FUS

For transient transfection experiments, neurons were fed DIV 6 and transfected with FLAG-HA-FUS constructs on DIV 7 using NeuroMag (Oz Biosciences NM51000, Marseille, France) transfection reagents. 1.0µg DNA and 1.75µL NeuroMag (for one 24-well well; 500uL volume) were combined in an eppendorf tube and brought up to a 50µL volume using neurobasal media. The DNA mixture was allowed to incubate for 20 minutes before addition to neurons. Upon addition, neuron cultures were placed on a NeuroMag magnet plate (Oz Biosciences MF10096) within the tissue culture incubator for 15 minutes to complete transfection. Transfected neurons were collected for experimental analyses on DIV 14-16.

### Lentiviral Production and Application

High titer lentivirus was prepared as described^69^. Briefly, HEK-293T cells were individually transfected using calcium phosphate with the shRNA or NLS-tdTomato-NES constructs described along with the packing plasmids: CMVdR8.91 plasmid and VSV-G. DNA constructs were prepared using by EndoFree Maxi Prep (Qiagen 12362, Germantown, MD, USA). Three hours after transfection, cell media was replaced with Opti-MEM (Invitrogen 31985070) and virus was collected in open-top Beckman tubes (Beckman Coulter 344058, Brea, CA, USA) by ultracentrifugation at 28,000 rpm in SW32Ti rotor 72 hours following transfection. Lentivirus titer was obtained by the transduction of HEK cells with serially diluted lentivirus. Upon titer determination, virus was added to DIV 6 non-cytosine β-D-arabinofuranoside hydrochloride treated neurons at an approximate titer of 1.2-1.8^10^ tu/ml. For all transduction experiments except fluorescence *in situ* hybridization, neurons were cytosine β-D-arabinofuranoside hydrochloride treated on DIV 7. Transduced neurons were collected 9 days post-transduction (DIV 15) for analysis.

### Immunofluorescence Analysis

Primary cortical and motor neurons were fixed with 4% paraformaldehyde (Fisher Scientific AAA1131336, Waltham, MA, USA) at room temperature for 15 minutes and permeabilized with 0.1-0.2% Triton X-100. Cortical neuron immunofluorescence experiments were conducted as described^12,26^ using antibodies listed in **Supplementary Table 1**. Primary motor neuron samples were blocked in 4% bovine serum albumin for 45 minutes and hybridized overnight at 4°C with primary antibodies (**Supplementary Table 1**) and AlexaFluor-conjugated secondary antibodies^68^.

### Image Acquisition and Processing

Primary motor neuron images were imaged using a widefield fluorescence microscope (Nikon TiE, Melville, NY, USA) equipped with a cooled CMOS camera (Andor, South Windsor, CT, USA). Primary motor neurons images were acquired as Z-stacks (0.2µm step size) using a 60x lens. As indicated, fixed primary cortical neurons were imaged using a Lecia TCS SP5 II laser scanning confocal (Leica Microsystems, Buffalo Grove, IL, USA) or Leica DMI6000B microscope as described^12^. For confocal images of whole cells, 12-bit stacks (Δz = 0.25µM steps, zoom = 3x, n = 23-30 planes) were acquired at 40x with a pixel size of 126nm (1024×1024 pixels; 1000Hz). For dendrites, 12-bit stacks (Δz = 0.08µM steps, zoom = 3x, n = 40-50 planes) were acquired at 63x using a pixel size of 80nm (1024×1024 pixels; 1000Hz). For fluorescence *in situ* hybridization (FISH), mFUS and somatic puromycin analyses, widefield stacks of the entire cell were acquired (z=0.2-.25µm) and deconvolved using the LAS AF One Software Blind algorithm (10 iterations). All neuron images were analyzed using MetaMorph software (Molecular Devices Inc., San Jose, CA, USA). The background and shading of stacks were corrected as described^26^. Sum or maximum projections were created from corrected stacks for downstream analyses.

For the quantification of cytoplasmic to nuclear (C:N) ratios, a 20×20 pixel region was applied to the nucleus and perinuclear area in the soma for each cell (visualized by DAPI and MAP2, NeuN or SMI-32 respectively) as well as an area within each image that contained no cells (representing background fluorescence). Using MetaMorph, the integrated intensity for the signal of interest was obtained for each region and a ratio of the cytoplasmic:nuclear signal was then generated following subtraction of background signal. For each experiment, the statistical comparison of C:N ratios with or without excitotoxic stress was completed using average C:N ratios collected from ≥three independent, biological experiments. For the analysis of FUS levels in neuronal dendrites, Microtubule-associated protein 2 (MAP2) was used visualize neuronal dendrites and create a dendritic mask using MetaMorph. Using the MAP2-defined mask, the integrated intensity of FUS staining was obtained and used to quantify the relative amount of FUS staining in dendrites. For the quantification of total neurons and neurons exhibiting FUS translocation, ≥10 fields of view were imaged at 40x for each condition tested. As indicated by MAP2 or meuronal nuclei (NeuN) staining, neurons were quantified from images with computer assistance from the ‘Cell Count’ feature in MetaMorph. To assess the percent neurons with protein translocation, cells were scored for the presence of cytoplasmic FUS and divided by the total neuron number to generate the percent population exhibiting a response.

### Puromycin Analysis

Based on previously described experiments^39^, 4mM stocks of puromycin (Invitrogen, A11138-03) were prepared in water. Neurons were treated with glutamate as described, except that a final concentration of 2µM puromycin was added to the primary cultured media during the last 15 minutes of the ‘washout’ period. As a positive control of translational inhibition, 100 μg/ml cycloheximide (MilliporeSigma C7698) throughout the experimental time course. Neurons were then analyzed by Western or Immunofluorescence using an anti-puromycin antibody (**Supplementary Table 1**). For the analysis of puromycin immunostaining upon FUS knockdown, a 20×20 pixel region was placed in the soma of GFP-positive cells. Using MetaMorph, the integrated intensity of this region was obtained and used to quantify relative puromycin levels as described.

### Fluorescence *in situ* Hybridization (FISH) Analysis

Non-cytosine β-D-arabinofuranoside hydrochloride treated neurons were plated on coverslips and transduced with shFUS or shSC-expressing lentivirus on DIV 6 and harvested on DIV 15. Following stress application, neurons were fixed with fresh 4% paraformaldehyde (Fisher Scientific F79-500) diluted in RNAse free water (Corning 46-000-CM) for 30 minutes at ambient temperature. FISH labeling was completed using a QuantiGene ViewRNA ISH Cell Assay Kit (Affymetrix QVC0001, Santa Clara, CA, USA) according to the manufacturer’s instructions. One exception to the protocol was that samples were dehydrated after fixation with two-minute incubations in 50%, 70%, and 100% ethanol at ambient temperature followed by a second addition of 100% ethanol and stored at −20°C for five days before processing. The Gria2-Cy3 probe was designed and tested for specificity by Affymetrix. For post-FISH immunofluorescence staining, after probe labeling, coverslips were washed in phosphate buffered saline for five minutes and then blocked and processed for immunofluorescence as described^70^. Coverslips were probed with MAP2 and GFP to visualize neurons and transduced cells, respectively. For analysis, neurons with at least 2 dendrites of 50+ µm lengths that did not excessively overlap with other cells were selected. Max projections of the imaged z-stacks were analyzed using MetaMorph software. For each neuron analyzed, 2-3 dendrites and the cell body were assessed for their area and the number of mRNA puncta present. Average dendrite data were reported for each cell and 10 cells were analyzed per construct/condition. Images were prepared for visualization in figures based upon methods previously described^71^.

### Lactate Dehydrogenase (LDH) Analysis

Neuron toxicity to glutamate was analyzed by the LDH assay using the CytoToxx 96 Non-Radioactive Cytotoxicity Assay kit (Promega G1782, Madison, WI, USA).

### Western Analysis

Neurons were treated, washed twice with phosphate buffered saline and lysed using RIPA buffer (Boston BioProducts BP-115-500, Ashland, MA, USA) supplemented with protease (Roche 11836170001, Basel, Switzerland) and phosphotase inhibitors (Roche 4906837001). Lysates were centrifuged at 13,500 rpm for 15 minutes, after which the supernatant was collected and its protein concentration determined using a bicinchoninic acid assay (Thermo Scientific Pierce 23227, Rockford, IL, USA). Lysates were subsequently used for Western and densitometry analysis as described^48^. Gels were loaded with 8-20µg lysate and GAPDH was used as a loading standard to determine relative protein levels. Primary antibodies used for analysis are described in **Supplementary Table 1**; LI-COR (Lincoln, NE, USA) secondary antibodies were used as described^48^.

## Results

### Excitotoxic levels of glutamate shift the nucleocytoplasmic equilibrium of disease-linked RNA binding proteins

To investigate a potential relationship between excitotoxicity and neurodegenerative disease-linked RBPs, we first examined whether excitotoxicity affects the nucleocytoplasmic equilibrium of a panel of proteins including FUS, TDP-43, hnRNP A1 and TATA-Binding Protein-Associated Factor 15 (TAF15). All four proteins have been linked to ALS^11^ and FUS, TDP-43 and TAF15 are also associated with FTD^27^. DIV 14–16 primary cortical neurons were bath treated with excitotoxic and physiologically relevant levels of glutamate^4,28^ (10µM; hereon referred to as Glu^excito^) for 10 minutes followed by a 30-minute washout period (Fig. 1A). Immunofluorescence was then used to assess the effect of Glu^excito^ on the cytoplasmic to nuclear (C:N) ratio of the endogenous RBPs (Fig. 1B-I). Strikingly, the FUS C:N ratio significantly increased ~15-fold from 0.04±0.05 to 0.6±0.3 in response to Glu^excito^ (Fig. 1B,F). This increase is likely due to a rapid egress of FUS from the nucleus into the cytoplasm, as a Western analysis revealed total FUS protein levels are unchanged before and after stress (**Fig. S1**). Glu^excito^ likewise induced a significant increase in the C:N ratio of TDP-43 (Fig. 1C,G) and hnRNPA1 (Fig. 1D,H) without altering protein expression (**Fig. S1**). Conversely, Glu^excito^ did not significantly alter the C:N ratio (Fig. 1E,I) or protein expression (**Fig. S1**) of TAF15.

**Figure 1.**
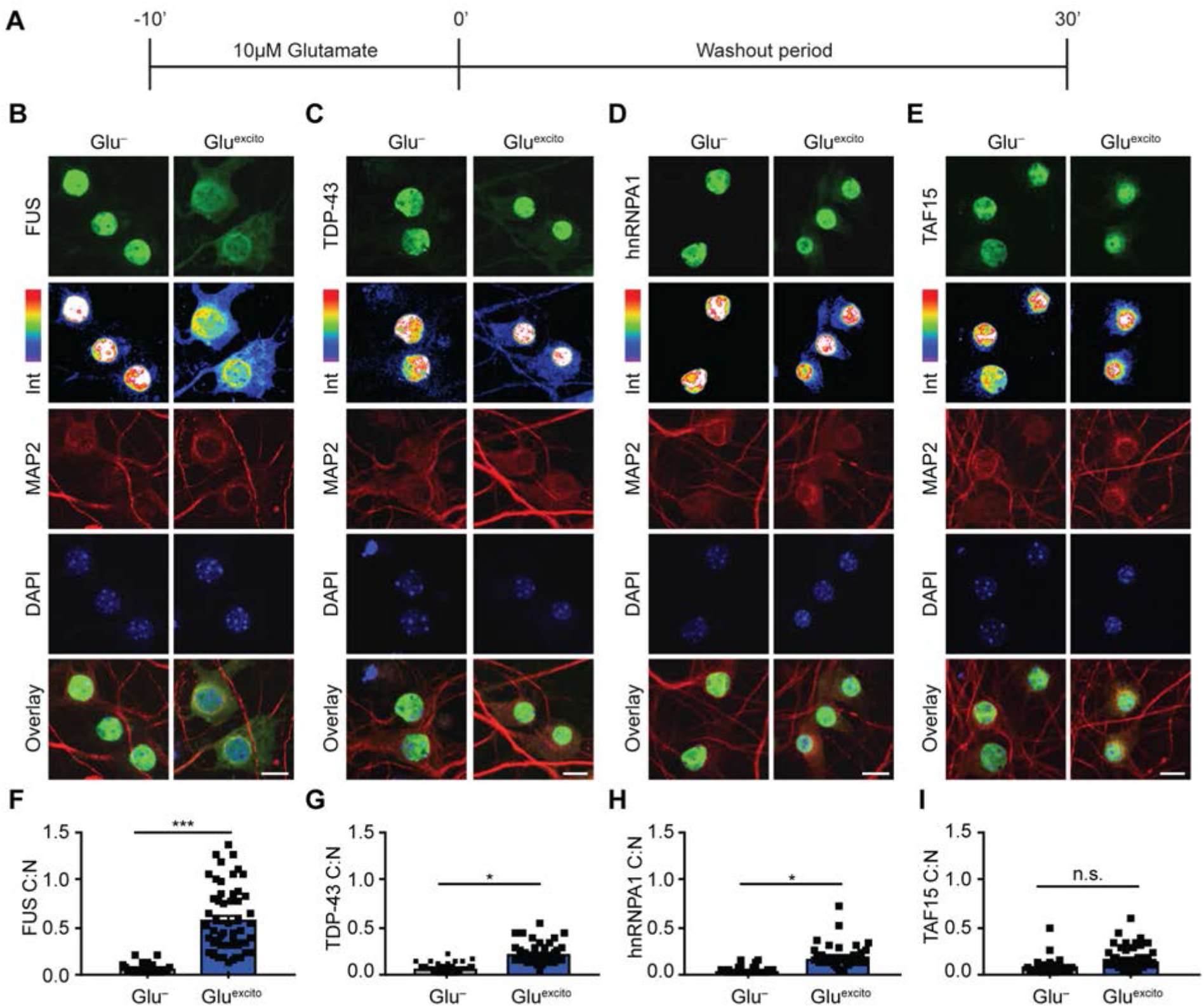
Endogenous FUS robustly translocates to the cytoplasm in response to excitotoxic stress. **(A)** DIV 14-16 primary cortical neurons were bath treated with 10µM glutamate (Glu^excito^) for 10 minutes, after which the glutamate-containing media was ‘washed out’ and replaced with cultured neuronal media for an additional 30 minutes. **(B-E)** Immunofluorescence and confocal microscopy revealed the cellular localization of FUS, TDP-43, hnRNPA1 and TAF15 (green) in the absence and presence of Glu^excito^. Endogenous RBP staining (green) visualized by a 16-color intensity map (Int) further demonstrates the cytoplasmic presence of these proteins. Neurons and dendrites were identified with anti-MAP2 staining (red), and nuclei with DAPI (blue). Scale bars = 10µm. **(F-I)** Quantification of the cytoplasmic to nuclear ratio (C:N) from (B-E). A significant nuclear egress of FUS (F), TDP-43 (G) and hnRNPA1 (H), but not TAF15 (I) was observed following Glu^excito^ treatment (n = 3-4 biological replicates). Black squares represent the C:N ratio of individual cells, and error bars correspond to SEM. Experimental means were calculated from the average C:N ratio across the individual biological replicates and significant comparisons were determined with a Student’s T-test (***p<0.001, *p<0.05, n.s. = non-significant).

In light of the robust response of FUS to Glu^excito^, we focused our attention on the properties of FUS translocation in more detail. First, endogenous FUS translocation in response to Glu^excito^ was confirmed using a panel of different anti-FUS antibodies (**Fig. S2 A,B**). We then examined the relationship between FUS translocation and glutamate concentration. With 10µM glutamate, the vast majority of neurons (91.3±11.5%) exhibited FUS egress (Fig. 2A,B), whereas 5% neurons exhibited translocation at ≤1μM, revealing a dependence of FUS localization on glutamate concentration (Fig. 2B). Within the time course of our experiment (Fig. 1A), a significant accumulation of endogenous FUS was also detected throughout MAP2-positive dendrites (Fig. 2C,D).

**Figure 2.**
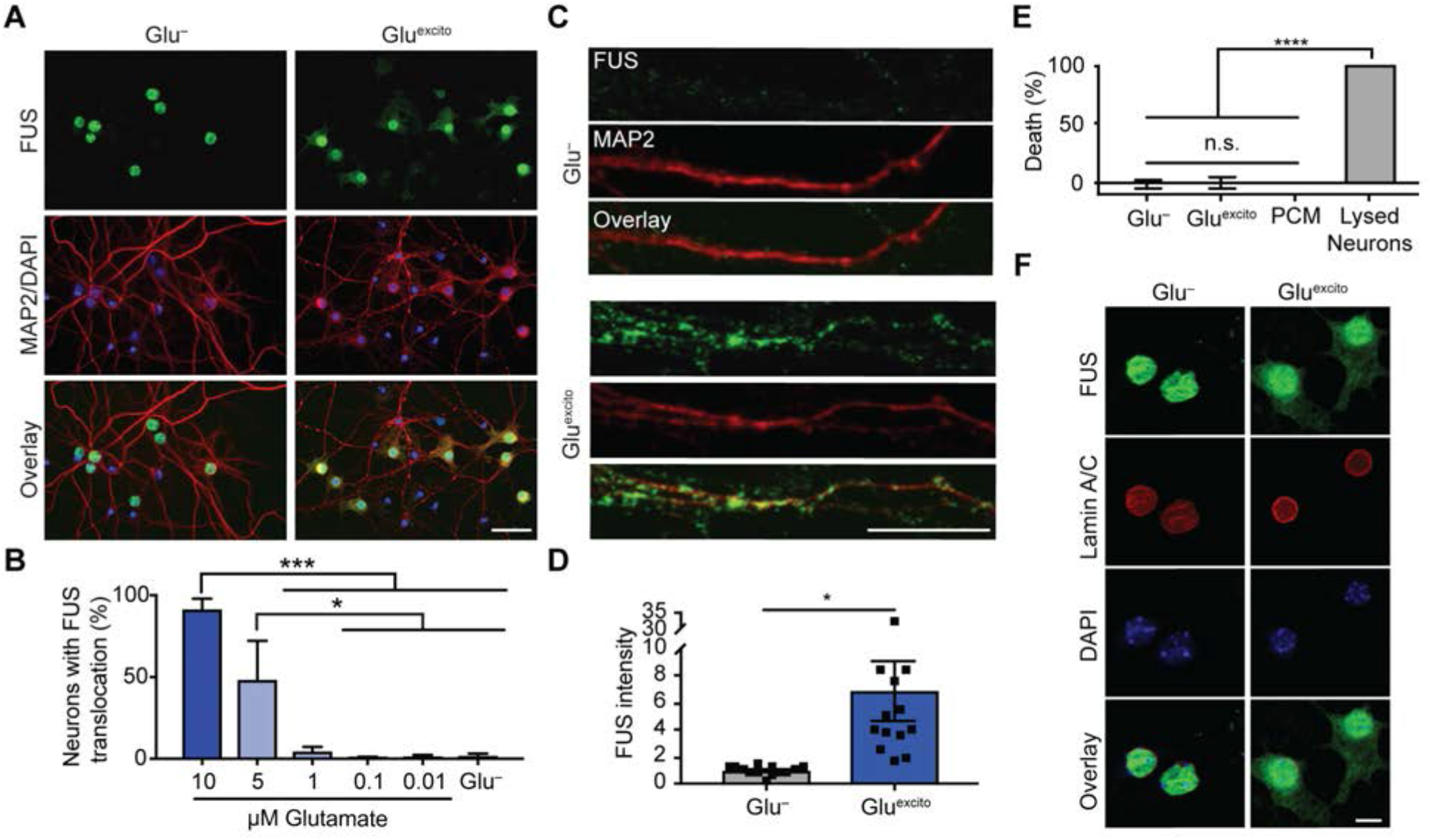
Cell viability and nuclear membrane integrity are intact under conditions of Glu^excito^ that promote FUS translocation. **(A)** Following excitotoxic insult, FUS egress and cytoskeletal rearrangements were detected by anti-FUS (green) and-MAP2 (red) staining, respectively. Scale bar = 40µm. **(B)** Quantification of (A) revealed a dependence of FUS translocation on the dose of glutamate in MAP2-positive neurons (one-way ANOVA and Tukey’s post-hoc test; ***p<0.001, *p<0.05; n = 3 biological replicates). **(C)** Increased dendritic FUS staining (green) was observed by confocal microscopy following excitotoxic stress. Dendrites were labeled with anti-MAP2 (red). Scale bar = 10µm. **(D)** Quantification of (C). Black squares represent the intensity of dendritic FUS staining per cell. Means represent the average of n = 4 biological replicates (Student’s T-test; *p<0.05) normalized to the control (Glu^−^). **(E)** Cytotoxicity induced by Glu^excito^ was assessed after the washout period (Fig. 1A) with the LDH assay. In contrast to the positive control (neurons treated with lysis buffer; lysed neurons), membrane permeabilization was not detected for neurons exposed to Glu^excito^. Neurons cultured in the absence of Glu^excito^ (Glu^−^) served as a negative control. Wells containing only primary neuron cultured medium (PCM) served as a background control. Results reflect n = 3 biological replicates analyzed with a one-way ANOVA and Tukey’s post-hoc test (****p<0.0001, n.s. = non-significant). (F) Immunofluorescence with anti-Lamin A/C staining (red) and confocal microscopy revealed the nuclear envelope was thickened yet still intact within neurons exhibiting translocated FUS (green) after Glu^excito^ exposure. The time point is the same as (E). Scale bar = 25µm. For (B), (D) and (E), error bars represent SEM.

Given the toxicity of Glu^excito^ on neurons^28^, we interrogated whether the rapid and robust accumulation of FUS outside the nucleus was simply a consequence of cell death and/or loss of nuclear envelope integrity. The extent of cell death was assessed using the LDH cytotoxicity assay, which detects the activity of LDH upon its release into the media from dead or dying cells. In contrast to neurons treated with lysis buffer, there was no evidence of cell death for neurons treated with Glu^excito^ (Fig. 2E). Further, Lamin A/C staining revealed an intact nuclear envelope in neurons exposed to Glu^excito^ (Fig. 2F). These observations support the premise that cytoplasmic FUS accumulation represents a cellular response to Glu^excito^, rather than a non-specific consequence of cell death. Moreover, RBP translocation appears selective, as TAF15 (Fig. 1E,I) and the cytoplasmic protein, fragile X mental retardation protein (FMRP; **Fig. S2C**), did not change localization following excitotoxic insult. It is noteworthy that Glu^excito^ affects neuron morphology at 30 minutes (Fig. 1A), potentially indicative of a stressed state. Anti-MAP2 staining revealed a rearrangement of the cytoskeleton; staining was more pronounced around the nucleus and indicated dendritic fragmentation (Fig. 1,2). Likewise, the nuclear lamina appeared thickened and the size of nuclei smaller in stressed neurons (Fig. 2F). As expected, neurons exposed to excitotoxic stimuli (10 μM, but not 1 µM glutamate) eventually undergo cell death within 24 hours of the initial insult^28^ (**Fig. S2D,E**).

### Excitotoxic stress induces egress of predominately nuclear ALS-linked FUS variants

The majority of ALS-linked mutations are located within the nuclear localization sequence (NLS) and, as such, these variants exhibit varying degrees of cytoplasmic mislocalization^15^. Given that both ALS-mutations and Glu^excito^ influence the subcellular localization of FUS, we investigated the relationship between these two factors. A series of FLAG-HA-tagged FUS variants were transiently expressed in neurons and the C:N ratio of exogenous FUS was determined in the absence and presence of Glu^excito^ (Fig. 3). In addition to wildtype (WT) FUS, we examined: H517Q, the only autosomal recessive FUS mutation associated with ALS^16^; R521G, representing a mutational ‘hot spot’ for ALS-FUS^29^; and R495X, a particularly aggressive ALS-linked mutation^26^. The degree of FUS mislocalization has been reported as H517Q≤R521G≪R495⨯ under basal conditions^26^, consistent with what was observed here (Fig. 3). As expected, FLAG-HA-FUS WT exhibited a significant translocation to the cytoplasm in response to Glu^excito^. Similarly, the C:N ratio of FLAG-HA-FUS H517Q and R521G, both of which exhibit a predominately nuclear localization under basal conditions (Fig. 3 and ^16,26^), also increased significantly with Glu^excito^. The C:N ratio of FLAG-HA-FUS R495X, which already exhibits a high degree of cytoplasmic localization under basal conditions (Fig. 3 and ^26^), did not change with Glu^excito^. This observation may be indicative of a ‘ceiling effect’, in that the normal nucleocytoplasmic distribution of R495X-FUS is equivalent to that of ‘maximally’ redistributed endogenous FUS following acute excitotoxic insult.

**Figure 3.**
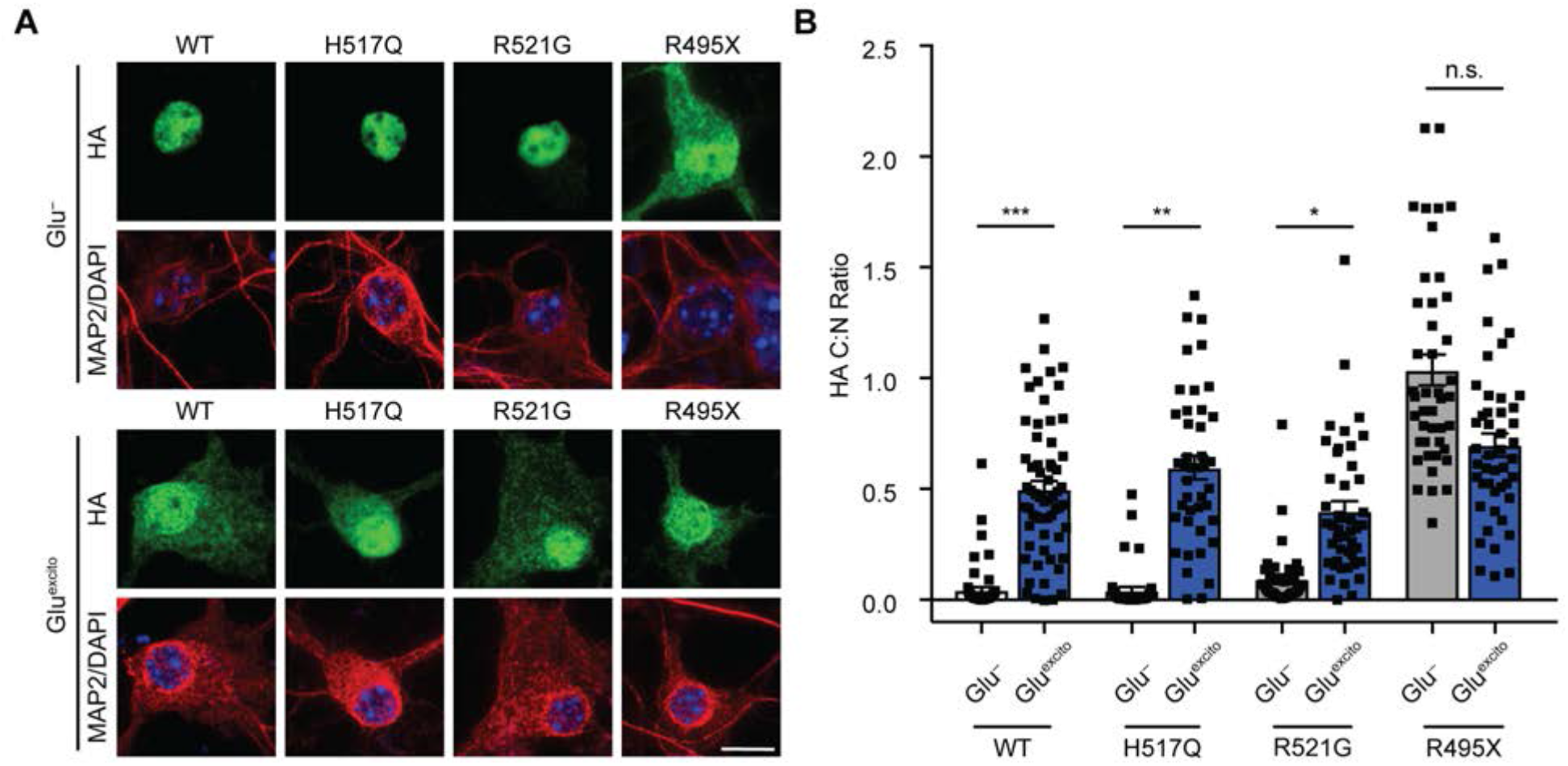
The effect of Glu^excito^ on ALS-linked FUS variants. **(A)** Cortical neurons transfected with the indicated FLAG-HA-tagged FUS variants were exposed to Glu^excito^ and nuclear FLAG-HA-FUS egress was assessed by immunofluorescence. Exogenous FUS was detected using an anti-HA antibody (green) within MAP2-positive neurons (red). Nuclei were stained with DAPI (blue). Scale bar = 10µm. **(B)** Quantification of the C:N ratio for FLAG-HA-FUS variants in (A) revealed a significant shift in equilibrium towards the cytoplasm for FLAG-HA-FUS WT, H517Q and R521G, but not R495X, in response to stress (Student’s T-test; ***p<0.001, *p<0.05, n.s. = not significant, n=3-5 biological experiments). Black squares represent individual, cellular C:N measurements. Error bars represent SEM.

### Nucleocytoplasmic transport is disrupted in response to excitotoxic stress

To understand the mechanism(s) underlying endogenous FUS egress in response to Glu^excito^, we began with an examination of nucleocytoplasmic transport factors. FUS contains two predicted chromosome region maintenance 1 (CRM1)-dependent nuclear export sequences (NES) within the RNA-recognition motif^30^. CRM1 is a major protein export factor, although whether this receptor controls nuclear FUS export is controversial^30,31^. To determine if excitotoxic FUS egress is CRM1-dependent, we pretreated neurons with the CRM1 inhibitor, KPT-330, prior to treatment with Glu^excito32^. The CRM1-dependent NLS-tdTomato-NES shuttling reporter was used as a positive control^33^. As expected, NLS-tdTomato-NES exhibited both a nuclear and cytoplasmic localization under basal conditions (Glu^−^,-KPT), whereas the localization of this reporter was significantly restricted to the nucleus in the presence of KPT-330 (Glu^−^, +KPT; Fig. 4A,B). In contrast, KPT-330 had no effect on nuclear FUS egress in response to Glu^excito^ (Glu^excito^ +/−KPT; Fig. 4A,C). Surprisingly, KPT-330 also failed to fully restrict NLS-tdTomato-NES to the nucleus under conditions of Glu^excito^ (Fig. 4A,B). Although there was a significant decrease in the percentage of cells with cytoplasmic NLS-tdTomato-NES in the presence of both KPT-330 and Glu^excito^ (60.1 ±8.0%) compared to Glu^excito^ alone (98.3±2.6, p=<0.0001), these results suggest that CRM1-mediated export is dysregulated under conditions of stress (Fig. 4B). Moreover, while endogenous CRM1 predominately localizes to the nucleus, Glu^excito^ induced a significant increase the number of neurons exhibiting a cytoplasmic localization of this protein (Fig. 4D,E). This finding prompted us to examine another critical nucleocytoplasmic transport factor, Ras-related nuclear protein (Ran). Ran is a GTPase that shuttles between the nucleus and cytoplasm and, depending on its nucleotide bound state, facilitates nuclear export or import^34^. Indeed, Glu^excito^ also induced a significant change in the nucleocytoplasmic distribution of Ran (Fig. 4F,G). Taken together, Glu^excito^ caused the redistribution of critical transport factors and attenuated the effects of KPT-330 on CRM1 export.

**Figure 4.**
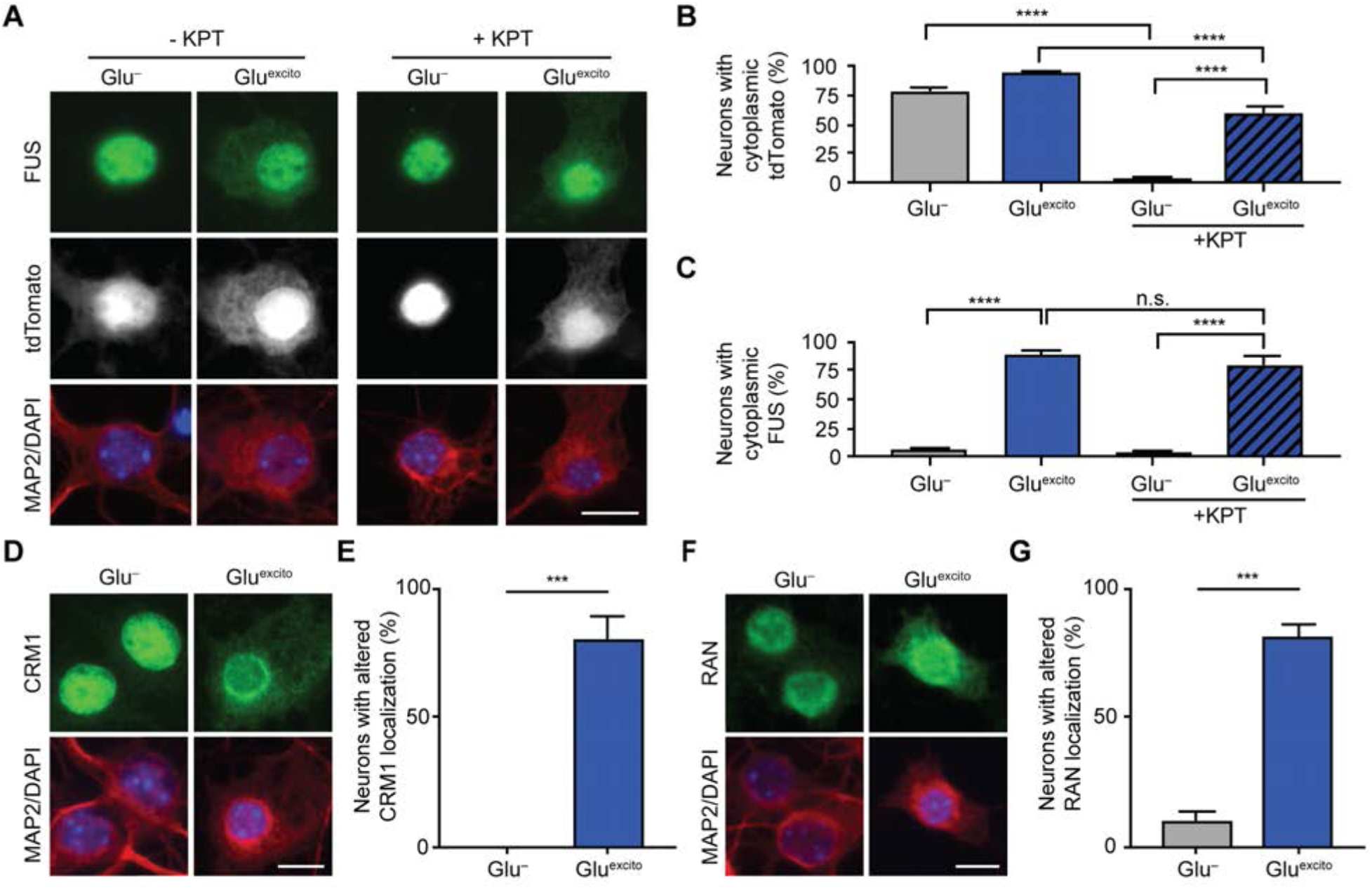
Nucleocytoplasmic transport is disrupted by Glu^excito^. **(A-C)** Cortical neurons expressing the shuttling reporter, NLS-tdTomato-NES, were treated with or without 500 nM of the exportin 1 inhibitor KPT-330 (KPT) prior to Glu^excito^ exposure. Neurons were identified with anti-MAP2 staining (A; red). The percentage of MAP2-positive cells expressing cytoplasmic NLS-tdTomato-NES (A; white) or FUS (A; green) was quantified in (B) and (C), respectively (n = 3 biological experiments). KPT-330 effectively prevents NLS-tdTomato-NES from localizing to the cytoplasm in the absence of stress (Glu-; A and B, two-way ANOVA and Tukey’s post-hoc test; ****p<0.0001), as expected. Conversely, in the presence of stress (Glu^excito^), KPT-330 fails to restrict NLS-tdTomato-NES and FUS localization to the nucleus, indicative of dysregulated nucleocytoplasmic transport (in B and C, compare Glu^−^ to Glu^excito^ in the presence of KPT-330, two-way ANOVA and Tukey’s post-hoc test; ****p<0.0001, n.s. = non-significant). The localization of nuclear transport factors CRM1 **(D, E)** and RAN **(F, G)** were significantly altered under conditions of Glu^excito^ in MAP2-positive neurons (red); CRM1 and RAN (green in D and E, respectively) were depleted from the nucleus (DAPI; blue) and exhibited a perinuclear accumulation. The percentage of neurons with CRM1 or RAN mislocalization were quantified in (E) and (G), respectively (Student’s T-test; ****p<0.0001, **p<0.01; n = 3 biological replicates). Error bars represent SEM. Scale bars = 10µm.

### Excitotoxic FUS egress is calcium dependent

Knowing that calcium influx is a critical component of excitotoxicity^1^, we sought to determine whether this signaling molecule is required for the response of FUS to excitotoxicity. To this end, the calcium chelator, EGTA, was included in the neuronal media during the experimental time course. Indeed, EGTA completely prevented Glu^excito^–induced FUS egress (Fig. 5A,B). Further application of the calcium ionophore, Ionomycin, was sufficient to induce FUS translocation in the vast majority (89.0±5.6%) of neurons (Fig. 5C,D). In light of our previous finding that hyperosmotic stress induces nuclear FUS egress^12^, we wondered whether calcium also mediated this response. In contrast to Glu^excito^, there was no effect of EGTA on FUS egress in neurons treated with hyperosmotic levels of sorbitol (Fig. 5E,F), indicative of distinct mechanisms for FUS egress under these stress conditions.

**Figure 5.**
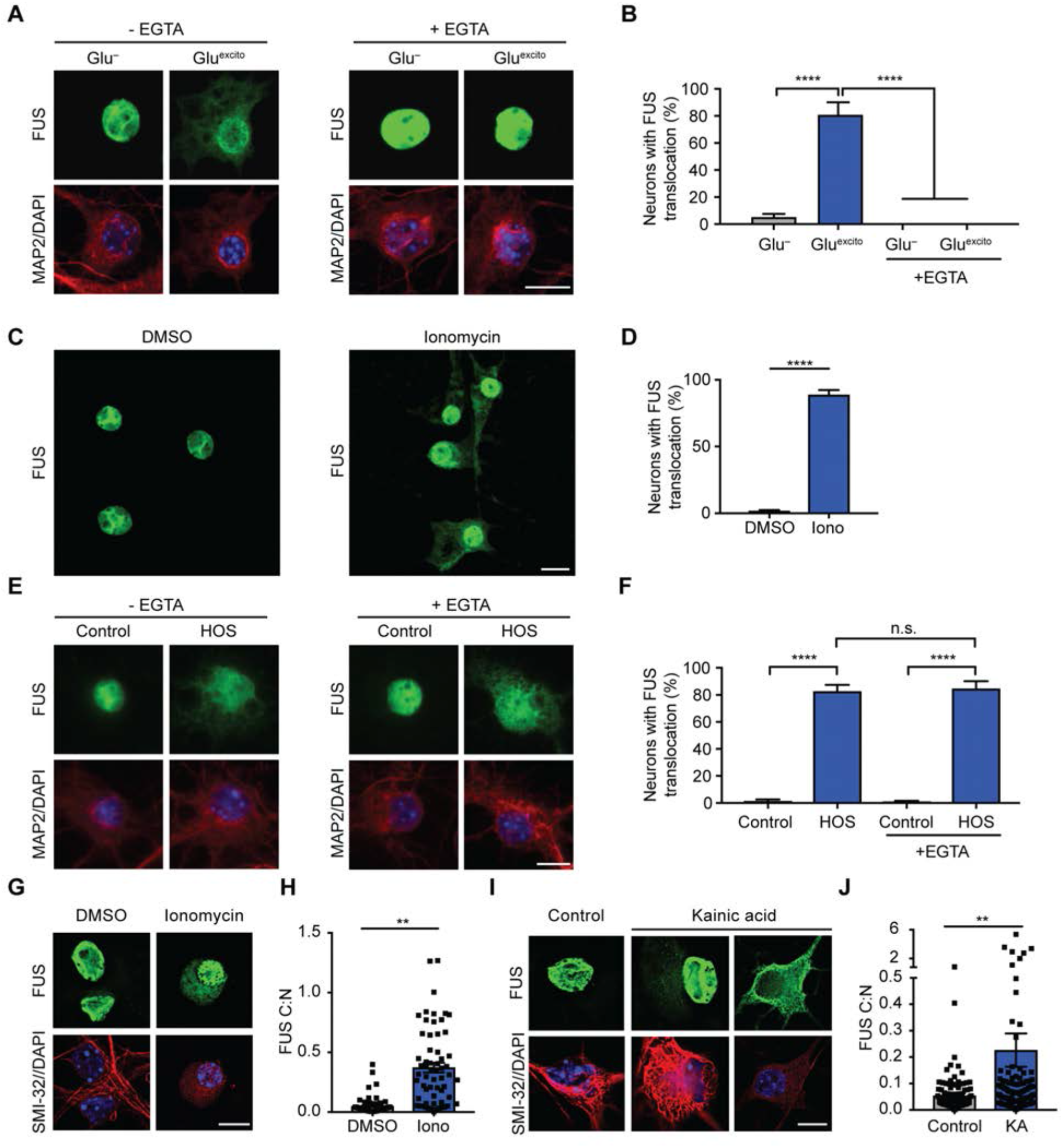
Calcium is necessary and sufficient for FUS translocation in primary cortical and motor neurons. **(A)** Reducing extracellular calcium levels with 2mM EGTA attenuates FUS egress (green) in MAP2-postive neurons (red) following excitotoxic insult. Nuclei were stained with DAPI (blue). **(B)** Quantification of confocal microscopy findings in (A) confirmed the effect of EGTA treatment (two-way ANOVA and Tukey’s post-hoc test; ****p<0.0001; n = 4 biological replicates). **(C,D)** Application of 10µM of the calcium ionophore, Ionomycin (Iono), for 1 hour induced FUS translocation relative to the dimethyl sulfoxide control (Student’s T-test; ****p<0.0001; n=3 biological replicates). **(E,F)** FUS translocation induced by hyperosmotic stress (HOS) was not significantly attenuated by EGTA treatment (two-way ANOVA and Tukey’s post-hoc test; ****p<0.0001, n.s. = non-significant; n=3 biological replicates). (**G,H**) Primary motor neurons treated with Ionomycin (Iono) as in (C,D) also exhibit FUS egress (green) and a significant increase in FUS C:N ratio (Student’s T-test, **p<0.01, n=3 biological replicates). Motor neurons were identified using the motor neuron marker, SMI-32 (red) and nuclei were stained with DAPI (blue). **(I)** A 10-minute treatment of 300µM kainic acid followed by a 1-hour recovery induced FUS egress in primary motor neurons. A near depletion of FUS from the nucleus was observed for a subset of motor neurons (kainic acid, *left*). **(J)** Kainic acid (KA) induced FUS egress was statistically significant relative to the washout control (Student’s T-test, **p<0.01, n=3 biological replicates). **(I, J)** Black squares indicate individual cell measurements normalized to the average of the replicate control. Accordingly, means represent the normalized average of n = 3 biological replicates. Error bars represent SEM. Scale bars = 10µm.

Next, we investigated the effect of calcium on FUS localization in primary motor neurons, the neuronal cell type predominately affected in ALS. Consistent with cortical neurons, the application of Ionomycin to DIV 6-8 motor neurons shifted the nucleocytoplasmic equilibrium of FUS towards the cytoplasm (Fig. 5G,H). Application of the glutamatergic agonist, kainic acid, to motor neurons also induced a significant increase in the C:N ratio of FUS (Fig. 5I,J). Kainic acid is known to induce motor neuron excitotoxicity^35^ and was used here to avoid confounding effects of glutamate uptake by astroglia present in the motor neuron cultures^36^. We noted a relatively wide range in the C:N ratio of FUS in kainic acid treated neurons; a sub-population of cells exhibited near complete egress of nuclear FUS (Fig. 5I,J), a result that was not observed in cortical neurons treated with glutamate.

### Excitotoxic stress induces translational repression independent of FUS expression and stress granule formation

Translational repression and stress granule formation are common cellular responses to stress^37,38^. Given that cytoplasmic forms of FUS have been linked to both translational regulation^39,40^ and stress granule formation^12,26^,^41,42^, we investigated both of these processes during excitotoxic stress. In contrast to neurons treated with sodium arsenite, a stressor known to induce the formation of stress granules^26^, Glu^excito^ did not induce the formation of Ras GTPase activating protein-binding protein 1 (G3BP1)-positive stress granules in neurons (**Fig. S3A**). Next, we assessed protein translation by pulse labeling neurons with puromycin, a small molecule that incorporates into elongating peptides^43^ (Fig. 6A). Detection of puromycin by immunofluorescence revealed a near-perfect correlation between neurons exhibiting FUS translocation and translational repression; all neurons with translocated FUS were puromycin-reduced, and vice versa (Fig. 6B). The degree of translational repression induced by Glu^excito^ was comparable to treatment with the translational inhibitor, cycloheximide (Fig. 6A-D), and did not promote FUS egress (Fig. 6B). Global translational repression was confirmed by a Western analysis (Fig. 6C, D), and was found to occur independently of eukaryotic translation initiation factor 2 alpha (EIF2a)-phosphorylation (**Fig. S3B,C**)^38^. However, endogenous FUS does not appear to play a vital role in regulating global translation, as puromycin levels were unaffected by FUS knockdown, both in the presence and absence of Glu^excito^ (**Fig. S3D-I**).

**Figure 6.**
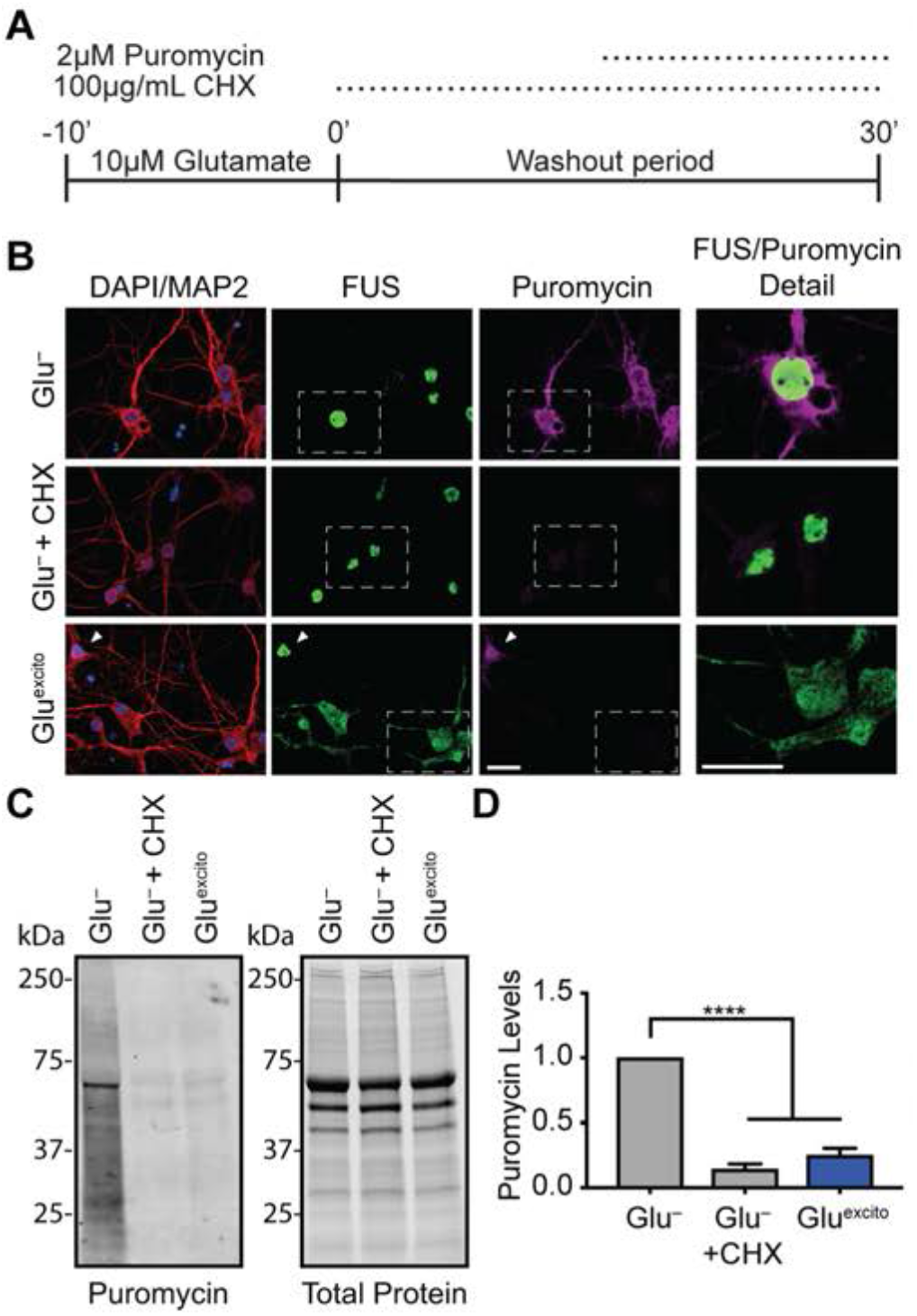
FUS translocation coincides with translational repression in neurons exposed to Glu^excito^. **(A)** Cellular translation in neurons was monitored by pulse-treatment and incorporation of the small molecule, puromycin, into nascent peptides during excitotoxic and/or cycloheximide treatment (CHX; inhibitor of protein translation). **(B)** The localization of FUS (green) and incorporated puromycin (magenta) in MAP2-postive neurons (red) was assessed by immunofluorescence. Relative to Glu^−^, protein translation was reduced upon application of cycloheximide or Glu^excito^, however cycloheximide did not induce FUS egress from nuclei (DAPI; blue). The white arrowhead marks a neuron with predominately nuclear FUS and high puromycin staining under Glu^excito^, whereas most neurons under this condition have cytoplasmic FUS and reduced puromycin staining. White boxes denote higher magnification details *(right)* to highlight neurons with representative levels of translation, as observed by anti-puromycin staining. **(C, D)** Western and densitometry analysis of puromycin incorporation confirms a significant reduction in translation following cycloheximide or Glu^excito^ treatment relative to Glu^−^ (one-way ANOVA and Tukey’s post-hoc test, ****p<0.0001, n = 3 biological replicates). Puromycin signal was normalized to total protein levels. Scale bars = 10µm. Error bars represent SEM.

### Gria2 mRNA is elevated in dendrites following excitotoxic insult in a FUS-dependent manner

RBPs such as FUS play crucial roles in mRNA processing^15^. Although FUS expression did not affect global protein synthesis (**Fig. S3E-I**), this analysis would not necessarily detect differences in the translation of specific transcripts, especially those targeted to dendrites for local translation^44^. Therefore, we investigated whether FUS modulates mRNA metabolism following excitotoxic insult and focused on Gria2, a transcript that is directly bound by FUS^45^. Gria2 mRNA encodes the GluR2 protein subunit of the AMPA receptor and has been implicated in calcium dyshomeostasis in both ALS^1^ and FTD^46^. Following depolarization, dendritic GluR2 expression is enhanced^47^. Under excitotoxic conditions, we uncovered a significant increase in Gria2 transcript density by FISH in both the soma and dendrites of cortical neurons (Fig. 7, **S4A-C**). To examine whether this increase in Gria2 mRNA density is FUS dependent, endogenous FUS levels were knocked down using two shRNAs targeting distinct sequences within FUS^48^ (**Fig. S3D-F**) prior to excitotoxic treatment (Fig. 7). Consistent with previous findings^45^, reduced FUS expression did not have a significant effect on the levels of Gria2 under basal conditions, as determined by FISH within the neuronal soma and dendrites (Fig. 7B,C). In contrast, Glu^excito^-induced changes to Gria2 were significantly attenuated upon FUS knockdown. Dendritic expression of Gria2 was particularly sensitive to FUS levels under Glu^excito^, as knockdown of FUS restored dendritic Gria2 levels to baseline (Fig. 7C, D). Within the time course of the analysis, we were unable to detect significant changes in GluR2 protein levels by Western blot analysis of whole cell lysates (**Fig. S4D,E**). Taken together, these data show that FUS expression is required for Glu^excito^-induced changes to Gria2 processing in neuronal dendrites (Fig. 8).

**Figure 7.**
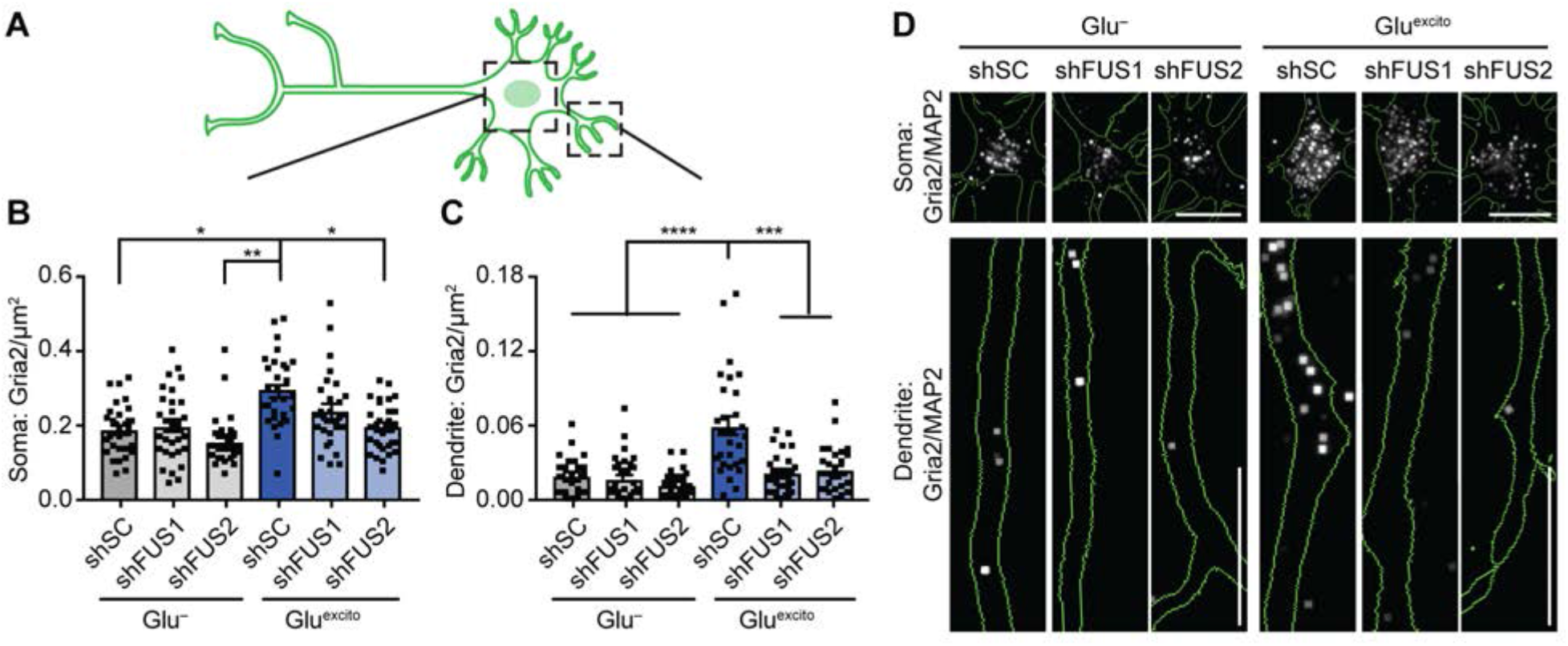
Elevation of Gria2 mRNA in dendrites following Glu^excito^ requires FUS expression. **(A)** Lentivirus expressing a GFP reporter and scrambled control shRNA (shSC) or shRNA against FUS (shFUS1, shFUS2) were used to reduce FUS levels in neurons in order to evaluate Gria2 mRNA distribution in soma and dendrites. **(B, C)** Following excitotoxic insult, the density of Gria2 was increased in both **(B)** soma and **(C)** dendrites of shSC transduced neurons; upon FUS knockdown dendritic Gria2 did not increase following treatment with Glu^excito^ (two-way ANOVA and Dunnett’s post-hoc test, ****p<0.0001, ***p<0.001, **p<0.01, *p<0.05, n = 3 biological replicates). Black squares indicate individual cell measurements. Error bars represent SEM. Scale bars = 10µm. **(D)** Representative images of (B,C). Gria2 mRNA was detected by FISH (white) in neurons outlined in green (raw images shown in Fig. S4B; image processing described in Fig. S4C).

**Figure 8.**
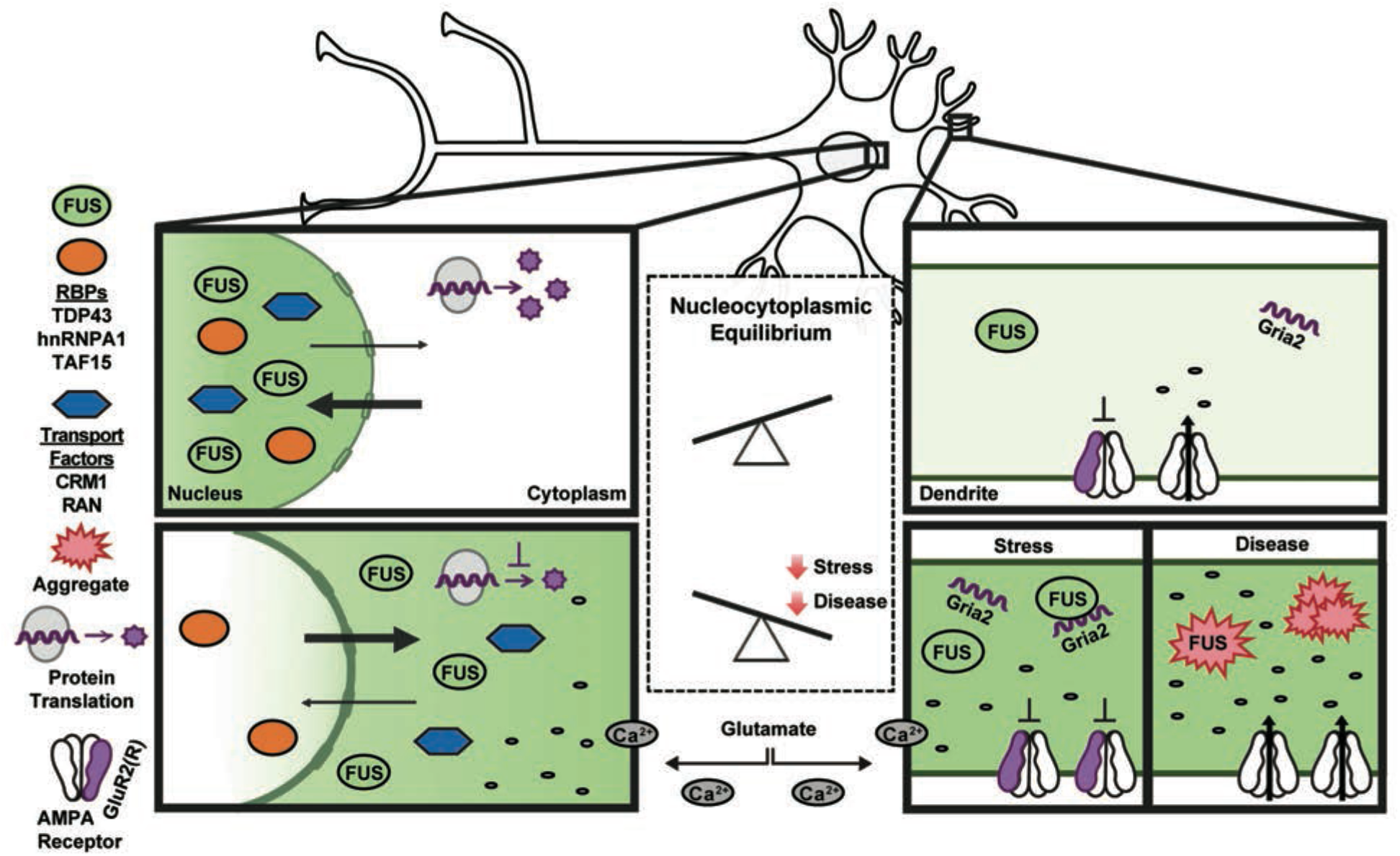
A model depicting the impact of excitotoxic stress on neuronal homeostasis and disease pathogenesis. Under homeostatic conditions, shuttling RBPs such as FUS are predominately localized within the nucleus (top). Excitotoxic levels of glutamate (bottom) induce a massive influx of calcium, which is sufficient to induce the robust nuclear egress of FUS into the neuronal soma and dendrites. Excitotoxic stress also leads to translational repression, a re-distribution of nucleocytoplasmic transport factors, and increased levels of Gria2 transcript within dendrites. The expression of FUS is required for enhanced levels of dendritic Gria2 in response to excitotoxic stress, implicating an RNA-processing role for FUS under these conditions. Enhanced levels of edited Gria2 transcript may represent a mechanism to offset the toxic effects of calcium influx. Prolonged or severe stress could manifest in the pathological aggregation of RBPs, including FUS, in neurodegenerative diseases such as ALS and FTD. Aberrant processing of Gria2 and/or GluR2 can occur through several mechanisms (e.g., expression of mutant FUS in astrocytes, loss of FUS function due to aggregation, and other means as described in the text), and contributes to calcium dyshomeostasis during disease.

## Discussion

This study uncovered an association between disease-linked RBPs and excitotoxicity, a stress that has particularly profound effects on the nucleocytoplasmic distribution of FUS in both cortical (Fig. 1, 2) and motor neurons (Fig. 5). There is a compelling body of evidence linking glutamate induced excitotoxicity to neurodegenerative diseases, including ALS^1–3^. For instance, elevated levels of glutamate were detected in biological samples from ALS patients^5,49,50^. Cell death caused by glutamate and calcium dysregulation has also been documented in multiple animal and cellular models^1,2^,5-^7,51^. The outcomes of this study shed new light on the excitotoxicity cascade and implicate, for the first time, a role for the ALS/FTD-linked protein FUS in this process.

Our results are consistent with a functional role for FUS in response to glutamatergic signaling^52^ rather than a non-specific effect of cell death. First, FUS egress precedes cell death (Fig. 2). Second, there is selectivity with respect to proteins that undergo a change in cellular localization; the response of FUS is particularly robust compared to the other proteins assessed in this study (Fig. 1, **S2C**, **S3A**). Third, the effects of excitotoxicity on Gria2 depend on FUS expression (Fig.7). FUS binds Gria2 mRNA within introns and the 3’ untranslated region, and Gria2 splicing is effected by FUS expression under basal conditons^45^. Under Glu^excito^, Gria2 density was enhanced in neuronal dendrites in a FUS-dependent manner (Fig. 7). Gria2 encodes the GluR2 protein subunit of the AMPA receptor. Normally, GluR2 is post-transcriptionally edited and GluR2-containing AMPA receptors are calcium impermeable. As such, the calcium permeability of AMPA receptors and the susceptibility of neurons to excitotoxicity is dependent on GluR2^1,8^. We speculate that the enhanced dendritic density of Gria2 may serve to increase the number of calcium impermeable (GluR2-containing) AMPA receptors and thereby offset calcium influx caused by existing calcium permeable (GluR2-lacking) receptors (Fig. 8). In ALS, this process could be compromised as a result of dysregulated Gria2 editing and/or GluR2 expression^8,53^, particularly in motor neurons that rely heavily on AMPA receptor signaling^1,2^. The effect of FUS on dendritic Gria2 density following Glu^excito^ (Fig. 7B,C) is novel and consistent with a role of FUS in modulating Gria2 processing. The exact nature of this role however remains to be fully elucidated, and could involve a function of FUS in Gria2 splicing^45^, transport^54^, and/or or stabilization^55,56^.

While investigating the mechanism(s) underlying excitotoxic FUS egress, we uncovered striking changes to the CRM1 nuclear export pathway (Fig. 4). Inhibition of CRM1-mediated export by KPT-330 failed to restrict both NLS-tdTomato-NES (Fig. 4A,B) and FUS (Fig. 4A,C) within the nucleus under Glu^excito^. Further, CRM1 localization was significantly shifted towards the cytoplasm (Fig. 4D,E). Despite these changes, nucleocytoplasmic transport was not completely dysregulated, as a partial inhibitory effect of KPT-330 on the shuttling reporter was observed (Fig. 4B). Our KPT-330 studies suggest that Glu^excito^-induced FUS egress occurs through a mechanism other than active CRM1 export, and could entail passive diffusion^30^ or alternative transport factors^57^. Selectivity of RBP egress following Glu^excito^ may stem from differences in nucleocytoplasmic shuttling dynamics, which are influenced by multiple factors including binding interactions and post-translational modifications^58^. An interesting area of future study could be to elucidate these factors and determine whether they are modulated by stress.

Alterations to CRM1 and Ran (Fig. 4) under Glu^excito^ may represent early signs of nucleocytoplasmic transport decline. Indeed, previous studies show that various forms of stress (e.g., excessive calcium influx, oxidative, and hyperosmotic stress) cause damage to nuclear pores and impair nucleocytoplasmic transport^59–63^. Mice deficient in key astroglial glutamate transporters exhibited both nuclear pore degradation and motor neuron degeneration^64^. Moreover, the nucleocytoplasmic transport pathway has been implicated in age-related neurodegeneration, particularly in the context of ALS and FTD^65^. While most ALS/FTD-associated studies have focused on the role of mutant proteins in dysregulating nucleocytoplasmic transport^34,65^, ALS/FTD-associated forms of cellular stress (e.g., excitotoxicity) may also contribute to nucleocytoplasmic transport defects in both inherited and sporadic forms of disease. In fact, nucleocytoplasmic transport is an emerging area of therapeutic development and the CRM1 inhibitor KPT-350 is advancing towards ALS clinical trials. Partial inhibition of CRM1 is expected to offset defects in nuclear import. CRM1 inhibitors have had a therapeutic effect in some^65,66^, but not all^57,64^, models of neurodegeneration. Collectively, the available data, including our own (Fig. 4), support CRM1-mediated nucleocytoplasmic transport as a viable therapeutic target for neurodegenerative disorders. However, a combination therapy addressing additional effects of stress-induced nuclear pore degradation (i.e., calpain inhibitors^64^) may be required for a significant therapeutic outcome.

The calcium-mediated response of FUS to Glu^excito^ has additional implications for neurodegeneration, including cases of FUS-mediated ALS. For instance, motor neurons derived from human ALS-FUS induced pluripotent stem cells are intrinsically hyperexcitable^67^. Further, the effects of ALS-linked FUS on calcium-mediated motor neuron toxicity is exacerbated by expression of the mutant protein in astrocytes^51,8^. Most ALS-linked FUS mutations are located within the NLS^29^ and induce a shift in the nucleocytoplasmic equilibrium of the protein toward the cytoplasm, where it is believed to exert a gain of toxic function^26,68^ (Fig. 3). As ALS-linked variants R521G and H517Q translocate further into the cytoplasm under Glu^excito^ (Fig. 3), we predict these and other variants with impaired binding to nuclear import factors will accumulate in the cytoplasm under conditions of chronic stress *in* vivo^41,59^. Moreover, chronic stress may result in nuclear depletion and cytoplasmic aggregation of wild-type FUS and TDP-43 in sporadic cases as well^19,20^. We propose a model whereby FUS and related RBPs play a functional role in response to normal stimulation and moderate degrees of stress, but that excessive or chronic stress severely disrupts their nucleocytoplasmic equilibrium and contributes to disease pathology (Fig.8).

## Acknowledgments

We are thankful to Drs. Kensuke Futai (University of Massachusetts Medical School; UMMS) and Miguel Sena-Esteves (UMMS) for sharing reagents and advice, Dr. Martin Hetzer (Salk Institute) for providing the NLS-tdTomato-NES construct and all the members of the Bosco and Landers labs for their valuable input. We are grateful to the following funding sources: US National Institutes of Health / National Institute on Neurological Disorders and Stroke R21NS091860 (DAB) and R01 NS078145 (DAB); ALS Association 18-IIA-418 (CF); Zelda Haidek Memorial Scholarship from UMMS (MT).

The authors declare no conflict of interest.

Supplementary information is available at *Cell Death and Differentiation’s* website.

---

**Supplementary Figure 1. RBP protein levels do not change in response to Glu^excito^**.
**(A, B)** Western analysis of cortical neurons demonstrate that FUS, TAF15, hnRNPA1 and TDP-43 protein levels do not change in response to Glu^excito^. **(C-F)** This observation was confirmed using densitometry. For quantification, RBP levels were first normalized to the loading standard, glyceraldehyde 3-phosphate dehydrogenase (GAPDH), and then the control condition, Glu^−^ (Student’s t-test, n.s. = non-significant, n=3 biological replicates). Error bars = SEM.

**Supplementary Figure 2.FMRP retains cytoplasmic localization following excitotoxic insult**.
**(A)** Anti-FUS antibody epitopes mapped to the domain structure of human FUS (QGSY = glycine-serine-tyrosine rich region, GLY = glycine-rich region, RRM = RNA recognition motif, RGG = arginine-glycine-glycine-rich region, ZF = zinc-finger domain and NLS = nuclear localization sequence). **(B)** Immunofluorescence staining of endogenous FUS (green) using antibodies with epitopes described in (A) consistently demonstrates FUS translocation following treatment with Glu^excito^. **(C)** Confocal analysis of anti-FMRP staining (green) demonstrates that the cytoplasmic localization of this protein is retained in neurons following excitotoxic stress (n = 2 biological replicates). Neurons were identified using a MAP2 antibody (red) and nuclei with DAPI (blue). **(D,E)** Quantification of MAP2-postive neurons at 24 hours relative to 30 minutes shows a significant reduction in neuron number following treatment with 10 but not 1µM glutamate relative to Glu^−^ (one-way ANOVA and Tukey’s post-hoc test, ***p<0.001, n.s. = non-significant, n = 3 biological replicates). Scale bars = 10µm. Error bars = SEM.

**Supplementary Figure 3. Reduced protein translation following excitotoxic stress is independent of EIF2a-phosphorylation and FUS levels**.
**(A)** Immunofluorescence staining of stress granule marker, G3BP1 (red), shows neurons treated sodium arsenite (NaAsO2) form stress granules unlike Glu^excito^ or Glu^−^ conditions where G3BP1 signal remains diffuse. Scale bar = 20µm. **(B,C)** Western and densitometry analysis demonstrate a significant increase in EIF2a phosphorylation (EIF2a-P) following sodium arsenite treatment (NaAsO2) relative to Glu^−^ but no significant change was observed for Glu^excito^. Levels of EIF2a-P were normalized to total EIF2a protein and the loading control, GAPDH (one-way ANOVA and Tukey’s post-hoc test, **p<0.01, n = 3 biological replicates). Scale bars = 10µm. Error bars represent SEM. **(D)** Primary neurons were transduced with shRNAs against mouse FUS (shFUS1, shFUS1) or a scrambled control (shSC) to induce FUS knockdown. Transduced neurons were identified by expression of a GFP reporter (white). Immunofluorescence staining of FUS (green) reveals FUS knockdown in transduced neurons identified using a MAP2 antibody (red). Scale bar = 50µm. **(E, F)** Western and densitometry analysis confirms FUS knockdown relative to non-transduced (NT) and shSC conditions. A modest increase in FUS levels was observed upon expression of shSC relative the loading standard, GAPDH (GAP; n=3; one-way ANOVA and Tukey’s Post Hoc test, ****p<0.0001, **p<0.01; n=3 biological replicates). **(G, H)** Neurons were pulse-chased labelled with puromycin (Puro; magenta) to assess nascent protein translation in transduced cells (as in B-D). The intensity of puromycin staining (Puro) for each condition was normalized to the respective stressed or unstressed non-transduced (NT) control. Scale bar = 10µm. **(I, J)** Quantification of puromycin (Puro) staining from (G,H) reveals no statistical difference in the somatic levels of translation following FUS knockdown (shFUS1, shFUS2) relative to shSC (one-way ANOVA and Dunnett’s post-hoc test, n.s. = not significant, n = 3 biological replicates). Error bars = SEM.

**Supplementary Figure 4. Steady state GluR2 protein levels are unchanged following Glu^excito^**.
**(A)**. Detection of the Gria2 transcript by FISH (green) was confirmed by the absence of signal in ‘no probe’ and ‘RNAse’ controls in MAP2-postive neurons (red). **(B)** Unprocessed images of Gria2 FISH in neurons represented in (Fig 7D). **(C)** To generated the images used in (Fig 7D), Gria2 puncta (green) were digitally dilated and coveted to white. Images of MAP2 staining used to indicate neurons and dendrites (red) were converted to binary and used to make a MAP2 mask subsequently outlined in green. White boxes exemplify approximate somatic (*) and dendritic (**) areas used for analysis and depiction in (Fig. 7). **(F, H)** Western and densitometry analysis of steady-state GluR2 protein levels reveal no statistical difference following Glu^excito^ relative to Glu^−^ and normalization to the loading standard, GAPDH (Student’s T-test, n.s. = not significant, n = 5 biological replicates). Scale bars = 25µm. Error bars = SEM.

## References

1 Van Den Bosch L, Van Damme P, Bogaert E, Robberecht W. The role of excitotoxicity in the pathogenesis of amyotrophic lateral sclerosis. Biochim Biophys Acta 2006; 1762: 1068–1082.

2 Starr A, Sattler R. Synaptic dysfunction and altered excitability in C9ORF72 ALS/FTD. Brain Research 2018; 1693: 98–108.

3 Fogarty MJ. Driven to decay: Excitability and synaptic abnormalities in amyotrophic lateral sclerosis. Brain Research Bulletin 2018; 140: 318–333.

4 Fiszman ML, Ricart KC, Latini A, Rodnguez G, Sica REP. In vitro neurotoxic properties and excitatory aminoacids concentration in the cerebrospinal fluid of amyotrophic lateral sclerosis patients. Relationship with the degree of certainty of disease diagnoses. Acta Neurol Scand 2010; 121: 120–126.

5 Spreux-Varoquaux O, Bensimon G, Lacomblez L, Salachas F, Pradat PF, Le Forestier N et al. Glutamate levels in cerebrospinal fluid in amyotrophic lateral sclerosis: a reappraisal using a new HPLC method with coulometric detection in a large cohort of patients. J Neurol Sci 2002; 193: 73–78.

6 Kawahara Yet al. Glutamate receptors: RNA editing and death of motor neurons. Nature 2004; 427: 801–801.

7 Hideyama Tet al. Profound downregulation of the RNA editing enzyme ADAR2 in ALS spinal motor neurons. Neurobiology of Disease 2012; 45: 1121–1128.

8 Van Damme Pet al. Astrocytes regulate GluR2 expression in motor neurons and their vulnerability to excitotoxicity. Proc Natl Acad Sci USA 2007; 104: 14825–14830.

9 Mitchell Jet al. Familial amyotrophic lateral sclerosis is associated with a mutation in D-amino acid oxidase. Proc Natl Acad Sci USA 2010; 107: 7556–7561.

10 Cheah BC, Vucic S, Krishnan AV, Kiernan MC. Riluzole, neuroprotection and amyotrophic lateral sclerosis. Curr Med Chem 2010; 17: 1942–1199.

11 Brown RH, Al-Chalabi A. Amyotrophic Lateral Sclerosis. N Engl J Med 2017; 377: 162–172.

12 Sama RRK et al. FUS/TLS assembles into stress granules and is a prosurvival factor during hyperosmolar stress. J Cell Physiol 2013; 228: 2222–2231.

13 Dewey CM et al. TDP-43 is directed to stress granules by sorbitol, a novel physiological osmotic and oxidative stressor. Mol Cell Biol 2011; 31: 1098–1108.

14 van Oordt WH et al. The MKK3/6-p38-signaling cascade alters the subcellular distribution of hnRNP A1 and modulates alternative splicing regulation. J Cell Biol 2000; 149: 307–316.

15 Sama R, Ward CL, Bosco DA.Functions of FUS/TLS From DNA Repair to Stress Response: Implications for ALS. ASN Neuro 2014; 6: 1–18.

16 Kwiatkowski TJ et al. Mutations in the FUS/TLS gene on chromosome 16 cause familial amyotrophic lateral sclerosis. Science 2009; 323: 1205–1208.

17 Vance Cet al. Mutations in FUS, an RNA processing protein, cause familial amyotrophic lateral sclerosis type 6. Science 2009; 323: 1208–1211.

18 Neumann Met al. A new subtype of frontotemporal lobar degeneration with FUS pathology. Brain 2009; 132: 2922–2931.

19 Keller BA, Volkening K, Droppelmann CA, Ang L-C, Rademakers R, Strong MJ. Co-aggregation of RNA binding proteins in ALS spinal motor neurons: evidence of a common pathogenic mechanism. Acta Neuropathol 2012; 124: 733–747.

20 Deng H-X et al. FUS-immunoreactive inclusions are a common feature in sporadic and non-SOD1 familial amyotrophic lateral sclerosis. Ann Neurol 2010; 67: 739–748.

21 Boyd JD et al. A high-content screen identifies novel compounds that inhibit stress-induced TDP-43 cellular aggregation and associated cytotoxicity. J Biomol Screen 2014; 19: 44–56.

22 Xu Get al. Identification of proteins sensitive to thermal stress in human neuroblastoma and glioma cell lines. PLoS ONE 2012; 7: e49021.

23 Kahl Aet al. Cerebral ischemia induces the aggregation of proteins linked to neurodegenerative diseases. Sci Rep 2018; 8: 2701.

24 Colombrita Cet al. TDP-43 is recruited to stress granules in conditions of oxidative insult. J Neurochem 2009; 111: 1051–1061.

25 McDonald KK et al. TAR DNA-binding protein 43 (TDP-43) regulates stress granule dynamics via differential regulation of G3BP and TIA-1. Hum Mol Gen 2011; 20: 1400–1410.

26 Bosco DA et al. Mutant FUS proteins that cause amyotrophic lateral sclerosis incorporate into stress granules. Hum Mol Gen 2010; 19: 4160–4175.

27 Ito D, Hatano M, Suzuki N. RNA binding proteins and the pathological cascade in ALS/FTD neurodegeneration. Sci Transl Med 2017; 9: eaah5436.

28 Schubert D, Piasecki D. Oxidative glutamate toxicity can be a component of the excitotoxicity cascade. J Neurosci 2001; 21: 7455–7462.

29 Lattante S, Rouleau GA, Kabashi E. TARDBP and FUS mutations associated with amyotrophic lateral sclerosis: summary and update. Hum Mutat 2013; 34: 812–826.

30 Ederle Het al. Nuclear egress of TDP-43 and FUS occurs independently of Exportin-1/CRM1. Sci Rep 2018; 8: 7084.

31 Kino Yet al. Intracellular localization and splicing regulation of FUS/TLS are variably affected by amyotrophic lateral sclerosis-linked mutations. Nucleic Acids Res 2011; 39: 2781–2798.

32 Grima JC, Daigle JG, Arbez N, Cunningham KC. Mutant Huntingtin Disrupts the Nuclear Pore Complex. Neuron 2017; 1: 93–107.

33 Hatch EM, Fischer AH, Deerinck TJ, Hetzer MW. Catastrophic nuclear envelope collapse in cancer cell micronuclei. Cell 2013; 154: 47–60.

34 Kim HJ, Taylor JP. Lost in Transportation: Nucleocytoplasmic Transport Defects in ALS and Other Neurodegenerative Diseases. Neuron 2017; 96: 285–297.

35 Fryer HJ, Knox RJ, Strittmatter SM, Kalb RG. Excitotoxic death of a subset of embryonic rat motor neurons in vitro. J Neurochem 1999; 72: 500–513.

36 Rose CR et al. Astroglial Glutamate Signaling and Uptake in the Hippocampus. Front Mol Neurosci 2017; 10: 451.

37 Kedersha N, Ivanov P, Anderson P. Stress granules and cell signaling: more than just a passing phase? Trends Biochem Sci 2013; 38: 494–506.

38 Holcik M, Sonenberg N. Translational control in stress and apoptosis. Nat Rev Mol Cell Biol 2005; 6: 318–327.

39 Murakami Tet al. ALS/FTD Mutation Induced Phase Transition of FUS Liquid Droplets and Reversible Hydrogels into Irreversible Hydrogels Impairs RNP Granule Function. Neuron 2015; 88: 678–690.

40 Yasuda Ket al. The RNA-binding protein Fus directs translation of localized mRNAs in APC-RNP granules. J Cell Biol 2013; 203: 737–746.

41 Dormann Det al. ALS-associated fused in sarcoma (FUS) mutations disrupt Transportin-mediated nuclear import. EMBO J 2010; 29: 2841–2857.

42 Gal Jet al. Nuclear localization sequence of FUS and induction of stress granules by ALS mutants. Neurobiol Aging 2011; 32: 2323.e27–40.

43 Schmidt EK, Clavarino G, Ceppi M, Pierre P. SUnSET, a nonradioactive method to monitor protein synthesis. Nat Methods 2009; 6: 275–277.

44 Holt CE, Schuman EM. The central dogma decentralized: new perspectives on RNA function and local translation in neurons. Neuron 2013; 80: 648–657.

45 Lagier-Tourenne C et al. Divergent roles of ALS-linked proteins FUS/TLS and TDP-43 intersect in processing long pre-mRNAs. Nat Neurosci 2012; 15: 1488–1497.

46 Gascon Eet al. Alterations in microRNA-124 and AMPA receptors contribute to social behavioral deficits in frontotemporal dementia. Nature Medicine 2014; 20: 1444–1451.

47 Ju Wet al. Activity-dependent regulation of dendritic synthesis and trafficking of AMPA receptors. Nat Neurosci 2004; 7: 244–253.

48 Ward CL et al. A loss of FUS/TLS function leads to impaired cellular proliferation. Cell Death Dis 2014; 5: e1572.

49 Plaitakis A, Constantakakis E. Altered metabolism of excitatory amino acids, N-acetyl-aspartate and N-acetyl-aspartylglutamate in amyotrophic lateral sclerosis. Brain Research Bulletin 1993.

50 Rothstein JD et al. Abnormal excitatory amino acid metabolism in amyotrophic lateral sclerosis. Ann Neurol 1990; 28: 18–25.

51 Kia A, McAvoy K, Krishnamurthy K, Trotti D, Pasinelli P. Astrocytes expressing ALS-linked mutant FUS induce motor neuron death through release of tumor necrosis factor-alpha. Glia 2018; 66: 1016–1033.

52 Fujii Ret al. The RNA binding protein TLS is translocated to dendritic spines by mGluR5 activation and regulates spine morphology. Curr Biol 2005; 15: 587–593.

53 Takuma H, Kwak S, Yoshizawa T, Kanazawa I. Reduction of GluR2 RNA editing, a molecular change that increases calcium influx through AMPA receptors, selective in the spinal ventral gray of patients with amyotrophic lateral sclerosis. Ann Neurol 1999; 46: 806–815.

54 Ling S-C. Synaptic Paths to Neurodegeneration: The Emerging Role of TDP-43 and FUS in Synaptic Functions. Neural Plast 2018; 2018: 8413496.

55 Udagawa Tet al. FUS regulates AMPA receptor function and FTLD/ALS-associated behaviour via GluA1 mRNA stabilization. Nat Commun 2015; 6: 7098.

56 Yokoi Set al. 3’UTR Length-Dependent Control of SynGAP Isoform α2 mRNA by FUS and ELAV-like Proteins Promotes Dendritic Spine Maturation and Cognitive Function. Cell Rep 2017; 20: 3071–3084.

57 Archbold HC et al. TDP43 nuclear export and neurodegeneration in models of amyotrophic lateral sclerosis and frontotemporal dementia. Sci Rep 2018; 8: 4606.

58 Rhoads SN, Monahan ZT, Yee DS, Shewmaker FP. The Role of Post-Translational Modifications on Prion-Like Aggregation and Liquid-Phase Separation of FUS. Int J Mol Sci 2018; 19. E886

59 Kodiha M, Chu A, Matusiewicz N, Stochaj U. Multiple mechanisms promote the inhibition of classical nuclear import upon exposure to severe oxidative stress. Cell Death Differ 2004; 11: 862–874.

60 Bano Det al. Alteration of the nuclear pore complex in Ca(2+)-mediated cell death. Cell Death Differ 2010; 17: 119–133.

61 Yasuda Y, Miyamoto Y, Saiwaki T, Yoneda Y. Mechanism of the stress-induced collapse of the Ran distribution. Exp Cell Res 2006; 312: 512–520.

62 Zhang Ket al. Stress Granule Assembly Disrupts Nucleocytoplasmic Transport. Cell 2018; 173: 958–971.e17

63 Kelley JB, Paschal BM. Hyperosmotic stress signaling to the nucleus disrupts the Ran gradient and the production of RanGTP. Mol Biol Cell 2007; 18: 4365–4376.

64 Sugiyama Ket al. Calpain-Dependent Degradation of Nucleoporins Contributes to Motor Neuron Death in a Mouse Model of Chronic Excitotoxicity. J Neurosci 2017; 37: 8830–8844.

65 Li N, Lagier-Tourenne C. Nuclear pores: the gate to neurodegeneration. Nat Neurosci 2018; 21: 156–158.

66 Haines JD et al. Nuclear export inhibitors avert progression in preclinical models of inflammatory demyelination. Nat Neurosci 2015; 18: 511–520.

67 Wainger BJ et al. Intrinsic Membrane Hyperexcitability of Amyotrophic Lateral Sclerosis Patient-Derived Motor Neurons. Cell Rep 2014; 7: 1–11.

68 Sama RRK et al. ALS-linked FUS exerts a gain of toxic function involving aberrant p38 MAPK activation. Sci Rep 2017; 7: 1205.

69 Sena-Esteves M, Tebbets JC, Steffens S, Crombleholme T, Flake AW. Optimized large-scale production of high titer lentivirus vector pseudotypes. J Virol Methods 2004; 122: 131–139.

70 Baron DM et al. Amyotrophic lateral sclerosis-linked FUS/TLS alters stress granule assembly and dynamics. Molecular Neurodegeneration 2013; 8: 30.

71 Cajigas IJ et al. The local transcriptome in the synaptic neuropil revealed by deep sequencing and high-resolution imaging. Neuron 2012; 74: 453–466.

